# Older mothers produce offspring with longer telomeres: a longitudinal within-parent analysis

**DOI:** 10.1101/2020.12.16.423019

**Authors:** Antony M. Brown, Emma M. Wood, Pablo Capilla-Lasheras, Xavier A. Harrison, Andrew J. Young

## Abstract

As telomere length often predicts survival and lifespan, there is considerable interest in understanding the origins of inter-individual variation in telomere length. Transgenerational effects of parental age on offspring telomere length are thought to be a key source of variation, but the rarity of longitudinal studies that examine the telomeres of successive offspring born throughout the lives of parents leaves such parental age effects poorly understood. Here, we exploit telomere length measures of successive offspring produced throughout the long breeding tenures of parents in wild white-browed sparrow-weaver (*Plocepasser mahali*) societies, to isolate the effects of within-parent changes in age on offspring telomere lengths. Our analyses reveal the first evidence to date of a positive within-parent effect of advancing age on offspring telomere length: as individual mothers age they produce offspring with longer telomeres. We consider the potential for pre- and post-natal mechanisms to explain our findings. As telomere erosion predicts offspring mortality in this species, this positive parental age effect could significantly impact parent and offspring fitness. Our findings support the view that transgenerational effects of parental age can be an appreciable source of inter-individual variation in telomere length.

## 1. Introduction

Telomeres are nucleoprotein complexes located at the ends of eukaryotic chromosomes [1]. They consist of a repeating non-coding DNA sequence (TTAGGGn in vertebrates) and have an important function in maintaining chromosomal integrity [1–3]. Telomeres vary in length both within and among individuals, and average telomere lengths shorten with advancing age in many taxa [2,3]. Telomeres may be shortened as a consequence of cell division and exposure to oxidative stress [4,5], and as critically short telomeres can trigger cellular senescence, excessive telomere shortening may have negative effects on performance [1–3,6–8]. Indeed, telomere length (TL) positively predicts survival and lifespan in a range of taxa (e.g. [6–8]) and is often considered a biomarker of somatic integrity [3]. Consequently, there is considerable interest in identifying the origins of the substantial inter-individual variation in TL within populations, given its potential implications for life-history trajectories and fitness. There is mounting evidence to suggest that parental age at offspring conception (hereafter ‘parental age’) predicts offspring TL in both human and non-human animals (e.g. [9–11]). However, the vast majority of studies to date have considered only whether population-level variation in parental age predicts offspring TL (see below). Whether such patterns reflect within-parent effects of advancing age rather than the selective disappearance of certain types of parents with advancing age, remains poorly understood.

In humans, population-level studies have consistently demonstrated that paternal (but not maternal) age at conception positively predicts offspring TL (e.g. [9,12]). In non-human animals however there is considerable variation among species in the nature of the population-level relationship between parental age and offspring TL. Studies in mammals, birds, and reptiles have to date found positive [13–15], negative [10,11,16], and no [17,18] relationship, with some finding evidence suggestive of either maternal [13,15] or paternal age effects [11,14,16], or both [10]. In order to understand the mechanistic origins and life-history implications of these relationships, studies now need to establish whether these patterns reflect within-parent effects of advancing age on the phenotypes of their offspring, or arise instead from among-parent processes (e.g. selective disappearance). Just two studies to date have yielded evidence that within-parent changes in age predict offspring TL (and see [19] for an example of no evident effects of within-parent changes in age on offspring TL). First, a study of captive zebra finches (*Taeniopygia guttata*) demonstrated that offspring produced by mothers when they were older (3.5 years of age) had shorter telomeres than those produced by the same mothers when young (6 months of age), while experimentally controlling paternal age [20]. Similarly, a study in a free-living population of jackdaws (*Corvus monedula*) found that as individual fathers age they produce offspring with shorter telomeres, while there was no effect of advancing maternal age [21]. These longitudinal studies therefore suggest that advancing parental age can *negatively* impact offspring TL.

Here, we investigate parental age effects on offspring TL in a free-living population of white-browed sparrow-weavers (*Plocepasser mahali*) in the Kalahari Desert. We use TL measures from successive offspring produced throughout the long breeding tenures of parents to isolate the effects of within-parent changes in age. White-browed sparrow-weavers live in year-round territorial groups, that comprise a single dominant male and female who completely monopolise within-group reproduction, and 0-12 non-breeding subordinate ‘helpers’ who assist with nestling feeding [22–24]. The rate of telomere attrition during development in this species predicts survival to adulthood [25], indicating that parental age effects on offspring TL could have substantial consequences for parent and offspring fitness.

## 2. Material and methods

### (a) Study species and field methods

Approximately 40 social groups have been monitored intensively since 2007 in Tswalu Kalahari Reserve (South Africa; 27°16’S, 22°25’E), leaving most individuals of known life-history (see [22–24]). Each group’s dominant breeding pair were identified using characteristic behavioural profiles, and were considered the parents of all offspring born in their group during their reproductive tenures (see electronic supplementary material [ESM] for justification and supporting analysis using genetic data). Nests were checked regularly throughout each breeding season (October to April inclusive) to determine egg lay and hatch dates, which allowed estimation of parental age (the parent’s age in days on the date that the focal offspring’s clutch was laid; see ESM) and offspring age at sampling (days since their hatch date). Blood samples for telomere assessments were routinely taken from nestlings and adult birds (see ESM). As helpers contribute to nestling feeding when present [22,23], we verified that our findings were unaffected by controlling for variation in an offspring’s rearing group size (see ESM).

### (b) Telomere length measurements

Telomere lengths were estimated from whole blood samples using qPCR analysis of relative telomere length (RTL) (following [26], details are described in ESM and [25]). We analysed 765 blood samples from 356 offspring from 248 clutches, produced by 61 mothers and 60 fathers in 41 social groups between 2010 and 2015 (numbers of mothers and fathers exceed the number of groups due to dominance turnovers during the study).

### (c) Statistical analyses

We fitted linear mixed-effects models with the lme4 package [27] in R (v3.6.1) [28] and ranked models by AICc (see ESM). We first investigated the effects of population-level variation in parental age on offspring RTL and, second, the effects of within-parent changes in age. In both cases, the following fixed effect predictors were fitted alongside parental ages: offspring sex and offspring age class (0-10 days [post-hatching], 11-89 days [later dependent period], and ≥90 days [independence and adulthood; mean ± SE age at sampling = 1.40 years ± 24.3 days; range 3 days – 7.35 years]). These offspring age classes were chosen to identify whether any parental age effects were present in offspring soon after hatching or arose later in offspring development (our work to date has revealed no evidence of changes in mean telomere length with advancing age in this population despite extensive longitudinal sampling; Wood et al. *in review*). Interactions between each parental age variable and offspring age were also included. Offspring ID, clutch ID nested within mother ID, qPCR plate ID, and sampling period (a factor identifying the breeding season and calendar year in which the blood sample was taken) were fitted as random intercepts. We did not fit father ID as it typically estimated zero or near-zero variance when alongside mother ID, due likely to its strong correlation with mother ID (dominant pairs can produce many clutches together; see ESM); our findings do not change if it is included. While global models contained both maternal and paternal age as fixed effects, models containing both were not AICc-ranked because maternal and paternal age were correlated (Pearson’s; r = 0.76, t = 31.97, df = 763, p < 0.001).

After conducting an initial analysis of the effects of population-level variation in maternal and paternal age on offspring RTL, we used within-subject centring [29] to partition the parental age variables into two components and repeated the analysis fitting both components as fixed effects: (i) the parent’s “mean age” across all sampled clutches for that parent and, (ii) the parent’s “Δ age”, the difference between the parental age value for the focal clutch and the parent’s mean age. The Δ age variables allow the models to estimate the effect of within-parent changes in age on offspring RTL, while the mean age variables estimate among-parent effects. As parents first encountered as fledglings or adults were assigned a minimum age (see ESM), any parental age relationships detected in the population-level analysis should be interpreted with caution. However, following within-subject centring, the Δ parental age values will be accurate for all parents.

## 3. Results

Analysis at the population level yielded support for a positive effect of maternal age on offspring RTL (Figure 1a; Table 1). Within-parent centring suggests that this pattern arises from a positive *within-mother* effect of advancing maternal age: as individual mothers get older they produce offspring with longer telomeres (Figure 1b; Table 2). The paternal age variables attracted consistently weaker support than maternal age variables (paternal age ΔAICc = +2.75; Table 1; paternal Δ age ΔAICc = +1.68; Table 2), suggesting that their positive relationships with offspring RTL could be a by-product of the strong correlation between maternal and paternal age. Our models did not yield clear evidence that the maternal age effect varies in strength with offspring age class (Table 1, 2). We confirmed that these parental age relationships cannot be attributed instead to correlated variation in social group size or uncertainty regarding parental age (see ESM).

**Figure 1.**
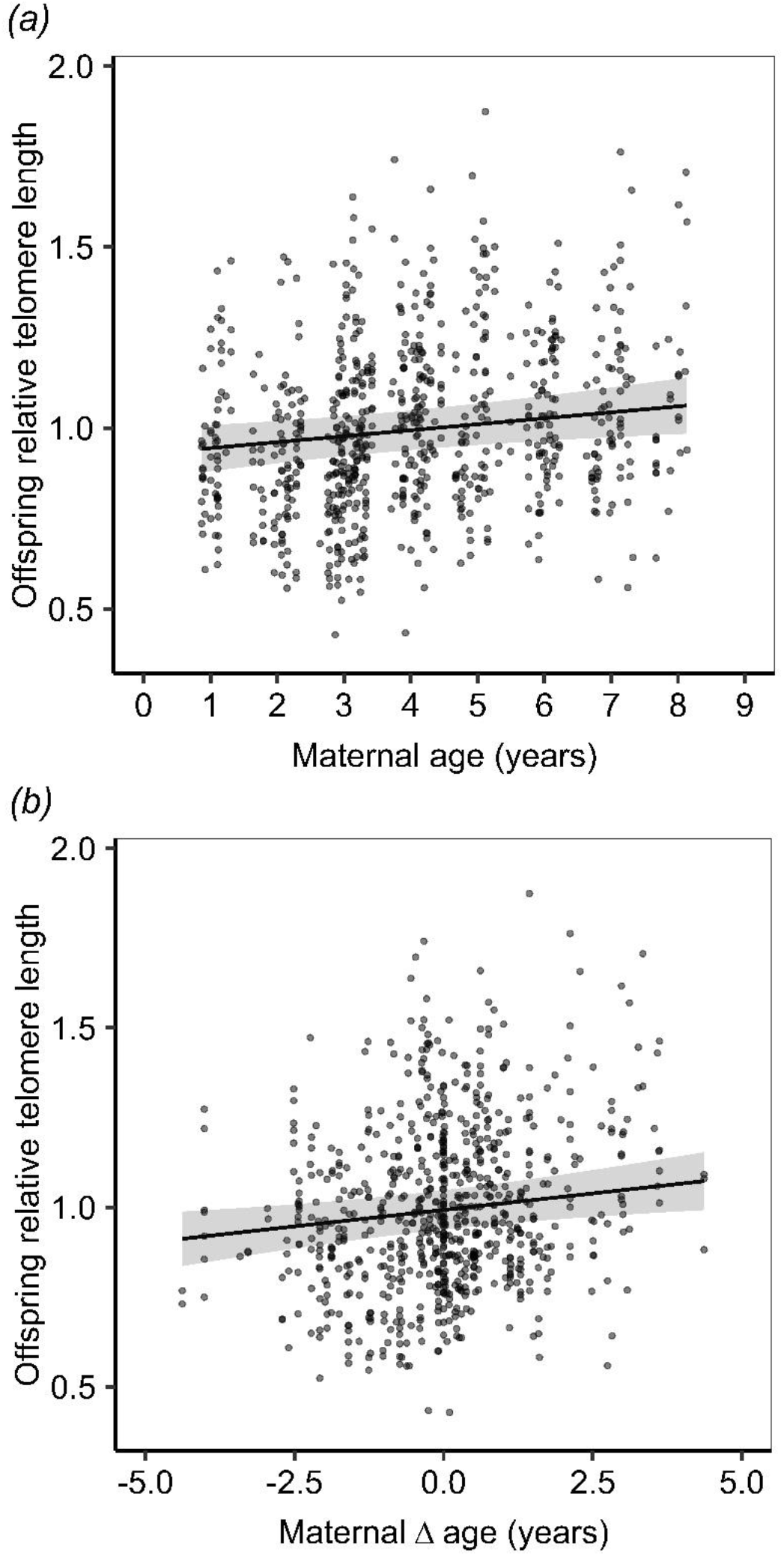
Relationship between maternal age and offspring RTL. Lines represent model predictions from the top-ranked models from our population-level *(a)* and within-parent *(b)* analyses. Shaded area represents upper and lower confidence intervals (1.96*SE). Points are unscaled raw data. There is some uncertainty in the maternal (and paternal) age values for some parents, while Δ age values are accurate for all birds (see methods).

**Table 1.** Δ6 AICc top model after implementation of the nesting rule [30] for the population-level analysis with scaled predictors of offspring RTL. Estimates are given with standard errors in parentheses.

| Intercept | Offspring<br>age class | Maternal<br>age | Paternal<br>age | Offspring age<br>class: paternal<br>age | df | logLik | AICc | Δ AICc | Adjusted<br>Akaike<br>weight |
| --- | --- | --- | --- | --- | --- | --- | --- | --- | --- |
| 0.995<br>(0.028) |  | 0.030<br>(0.011) |  |  | 8 | 253.04 | -489.9 | 0.00 | 0.634 |
| 0.970<br>(0.033) | + |  | + | + | 12 | 255.78 | -487.1 | 2.75 | 0.160 |
| 0.989<br>(0.029) |  |  | 0.023<br>(0.013) |  | 8 | 251.42 | -486.7 | 3.24 | 0.126 |
| 0.988<br>(0.030) |  |  |  |  | 7 | 249.95 | -485.7 | 4.14 | 0.080 |

Predictors absent from top model set: Offspring sex, Offspring age class: maternal age interaction. Paternal
age effect estimates in second-ranked model: for offspring age 0-10 days = -0.017 (0.021); 11-89 days = 0.0003
(0.0178); ≥90 days = 0.037 (0.015).

**Table 2.**
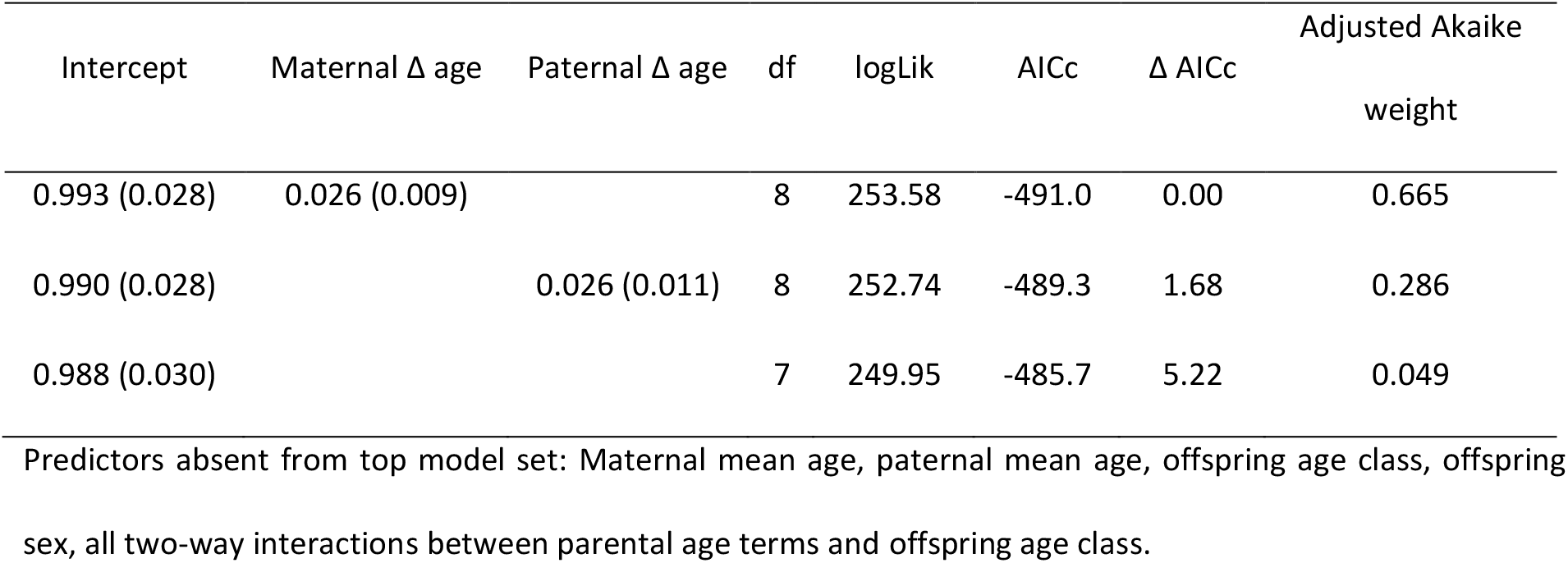
Δ6 AICc top model set after implementation of the nesting rule [30] for the *within-parent* analysis with scaled predictors of offspring RTL. Estimates are given with standard errors in parentheses.

| Intercept | Maternal $\Delta$ age | Paternal $\Delta$ age | df | logLik | AICc | $\Delta$ AICc | Adjusted Akaike weight |
| --- | --- | --- | --- | --- | --- | --- | --- |
| 0.993 (0.028) | 0.026 (0.009) |  | 8 | 253.58 | -491.0 | 0.00 | 0.665 |
| 0.990 (0.028) |  | 0.026 (0.011) | 8 | 252.74 | -489.3 | 1.68 | 0.286 |
| 0.988 (0.030) |  |  | 7 | 249.95 | -485.7 | 5.22 | 0.049 |

Predictors absent from top model set: Maternal mean age, paternal mean age, offspring age class, offspring sex, all two-way interactions between parental age terms and offspring age class.

## 4. Discussion

Our analyses reveal that maternal age positively predicts offspring TL at the population level and suggest that this relationship reflects positive effects of *within-mother* changes in maternal age rather than confounding effects of variation among mothers. A within-parent effect of parental age on offspring TL has only been reported in two other studies (also in birds), both of which reported a negative effect [20,21]. Our findings therefore constitute the first evidence of a positive within-parent effect of advancing parental age on offspring TL.

Parental age effects have the potential to arise via both pre-natal mechanisms (e.g. gamete-mediated epigenetic effects) and post-natal mechanisms (including age-related changes in the provision of post-natal care), as has recently been underscored by evidence of both biological parent and foster parent age effects on offspring TL [10,21]. Our models did not yield clear statistical support for an interactive effect of maternal age and offspring age on offspring TL, despite large samples sizes in each offspring developmental age class (see ESM). This highlights the possibility that this effect arose via *pre-natal* mechanisms, such as maternal modification of egg constituents with advancing age [31]. However, there is a need for caution in discounting a role for post-natal mechanisms, as the maternal age effect size quadruples between early development and the period of independence and adulthood (Figure S1). A number of mechanisms exist by which a relationship could arise post-natally. For example, the quality of post-natal maternal care could conceivably increase as mothers age, leading to slower rates of telomere attrition during the nestling period [32]. Such a pattern could reflect beneficial effects on offspring of increasing maternal breeding experience (e.g. [33]). While negative effects of parental age (e.g. [10,21]) might be expected to manifest as parents senesce, the senescent period may not be well represented within our data as sampled mothers were not known to exceed 8 years of age and the species can breed beyond 12 years of age [25].

Offspring produced by older parents often have reduced fitness and lifespan (the ‘Lansing effect’ [34,35]). By contrast, our finding of a positive effect of maternal age on offspring TL highlights the possibility in this species of parental age-related increases in offspring performance, given that offspring TL often positively predicts their downstream performance [25]. Indeed, recent work on another social vertebrate has yielded striking evidence of a positive maternal age effect on offspring performance [36]. These apparent benefits of advanced parental age highlight the wider need for analyses of the fitness implications of reproductive senescence to consider parental age-related changes in offspring quality as well as offspring production, given the potential for the former to partially offset the latter.

To conclude, we present the first evidence that within-parent changes in age positively predict offspring TL. Our findings, coupled with those of other recent studies that have demonstrated within-parent effects of advancing age on offspring TL [20,21], suggest parental age at reproduction does partially explain inter-individual variation in TL in animal populations. Indeed, follow-up analyses reveal that this maternal age effect in white-browed sparrow-weaver societies persists into offspring adulthood (see ESM). Work is now needed to elucidate the mechanisms driving these parental age effects and to establish the causes of their variation across taxa.

## Supporting information

Electronic Supplementary Material

## Ethics

The protocols followed in this study were approved by University of Pretoria Animal Ethics Committee (EC023-07 and EC100-12).

## Data accessibility

Data are available on Dryad (doi:10.5061/dryad.2jm63xsng) [37].

## Authors’ contributions

A.M.B., E.M.W. and A.J.Y. designed the study. E.M.W. conducted the fieldwork and performed the telomere length analyses. A.J.Y. led the long-term field study. A.M.B., P.C-L. and X.A.H. conducted the microsatellite genotyping analyses. A.M.B. analysed the data and wrote the manuscript. All authors commented on the manuscript and approved the final version.

## Acknowledgements

We thank the many sparrow-weaver team members for their contributions to the collection of our long-term life history data, Nigel Bennett for logistical support, Northern Cape Conservation for permission to carry out the research, and E. Oppenheimer & Son, the Tswalu Foundation, and all at Tswalu Kalahari Reserve for their support in the field.

## References

1. Blackburn EH. 2001 Switching and signaling at the telomere. Cell 106, 661–673. (doi:10.1016/S0092-8674(01)00492-5)

2. Monaghan P. 2010 Telomeres and life histories: the long and the short of it. Ann. N. Y. Acad. Sci. 1206, 130–142. (doi:10.1111/j.1749-6632.2010.05705.x)

3. Young AJ. 2018 The role of telomeres in the mechanisms and evolution of life-history trade-offs and ageing. Phil. Trans. R. Soc. B 373, 20160452. (doi:10.1098/rstb.2016.0452)

4. Monaghan P, Ozanne SE. 2018 Somatic growth and telomere dynamics in vertebrates: relationships, mechanisms and consequences. Phil. Trans. R. Soc. B 373, 20160446. (doi:10.1098/rstb.2016.0446)

5. von Zglinicki T. 2002 Oxidative stress shortens telomeres. Trends Biochem. Sci. 27, 339–344. (doi:10.1016/S0968-0004(02)02110-2)

6. Heidinger BJ, Blount JD, Boner W, Griffiths K, Metcalfe NB, Monaghan P. 2012 Telomere length in early life predicts lifespan. Proc. Natl. Acad. Sci. U. S. A. 109, 1743–1748. (doi:10.1073/pnas.1113306109)

7. Bize P, Criscuolo F, Metcalfe NB, Nasir L, Monaghan P. 2009 Telomere dynamics rather than age predict life expectancy in the wild. Proc. R. Soc. B 276, 1679–1683. (doi:10.1098/rspb.2008.1817)

8. Wilbourn RV, Moatt JP, Froy H, Walling CA, Nussey DH, Boonekamp JJ. 2018 The relationship between telomere length and mortality risk in non-model vertebrate systems: a meta-analysis. Phil. Trans. R. Soc. B 373, 20160447. (doi:10.1098/rstb.2016.0447)

9. Eisenberg DTA, Hayes MG, Kuzawa CW. 2012 Delayed paternal age of reproduction in humans is associated with longer telomeres across two generations of descendants. Proc. Natl. Acad. Sci. U. S. A. 109, 10251–10256. (doi:10.1073/pnas.1202092109)

10. Criscuolo F, Zahn S, Bize P. 2017 Offspring telomere length in the long lived Alpine swift is negatively related to the age of their biological father and foster mother. Biol. Lett. 13, 20170188. (doi:10.1098/rsbl.2017.0188)

11. Olsson M, Pauliny A, Wapstra E, Uller T, Schwartz T, Blomqvist D. 2011 Sex differences in sand lizard telomere inheritance: paternal epigenetic effects increases telomere heritability and offspring survival. PLoS One 6, e17473. (doi:10.1371/journal.pone.0017473)

12. Unryn BM, Cook LS, Riabowol KT. 2005 Paternal age is positively linked to telomere length of children. Aging Cell 4, 97–101. (doi:10.1111/j.1474-9728.2005.00144.x)

13. Cram DL, Monaghan P, Gillespie R, Clutton-Brock T. 2017 Effects of early-life competition and maternal nutrition on telomere lengths in wild meerkats. Proc. R. Soc. B 284, 20171383. (doi:10.1098/rspb.2017.1383)

14. Eisenberg DTA, Tackney J, Cawthon RM, Cloutier CT, Hawkes K. 2016 Paternal and grandpaternal ages at conception and descendant telomere lengths in chimpanzees and humans. Am. J. Phys. Anthropol. 162, 201–207. (doi:10.1002/ajpa.23109)

15. Asghar M, Bensch S, Tarka M, Hansson B, Hasselquist D. 2015 Maternal and genetic factors determine early life telomere length. Proc. R. Soc. B 282, 20142263. (doi:10.1098/rspb.2014.2263)

16. Bouwhuis S, Verhulst S, Bauch C, Vedder O. 2018 Reduced telomere length in offspring of old fathers in a long-lived seabird. Biol. Lett. 14, 20180213. (doi:10.1098/rsbl.2018.0213)

17. Froy H et al. 2017 No evidence for parental age effects on offspring leukocyte telomere length in free-living Soay sheep. Sci. Rep. 7, 9991. (doi:10.1038/s41598-017-09861-3)

18. Belmaker A, Hallinger KK, Glynn RA, Winkler DW, Haussmann MF. 2019 The environmental and genetic determinants of chick telomere length in Tree Swallows (*Tachycineta bicolor*). Ecol. Evol. 9, 8175–8186. (doi:10.1002/ece3.5386)

19. van Lieshout SHJ, Sparks AM, Bretman A, Newman C, Buesching CD, Burke T, Macdonald DW, Dugdale HL. 2020 Estimation of environmental, genetic and parental age at conception effects on telomere length in a wild mammal. J. Evol. Biol. 00, 1–13. (doi:10.1111/jeb.13728)

20. Marasco V, Boner W, Griffiths K, Heidinger B, Monaghan P. 2019 Intergenerational effects on offspring telomere length: interactions among maternal age, stress exposure and offspring sex. Proc. R. Soc. B 286, 20191845. (doi:10.1098/rspb.2019.1845)

21. Bauch C, Boonekamp JJ, Korsten P, Mulder E, Verhulst S. 2019 Epigenetic inheritance of telomere length in wild birds. PLoS Genet. 15, e1007827. (doi:10.1371/journal.pgen.1007827)

22. Harrison XA, York JE, Cram DL, Young AJ. 2013 Extra-group mating increases inbreeding risk in a cooperatively breeding bird. Mol. Ecol. 22, 5700–5715. (doi:10.1111/mec.12505)

23. Harrison XA, York JE, Cram DL, Hares MC, Young AJ. 2013 Complete reproductive skew within white-browed sparrow weaver groups despite outbreeding opportunities for subordinates of both sexes. Behav. Ecol. Sociobiol. 67, 1915–1929. (doi:10.1007/s00265-013-1599-1)

24. Cram DL, Blount JD, Young AJ. 2015 The oxidative costs of reproduction are group-size dependent in a wild cooperative breeder. Proc. R. Soc. B 282, 20152031. (doi:10.1098/rspb.2015.2031)

25. Wood EM, Young AJ. 2019 Telomere attrition predicts reduced survival in a wild social bird, but short telomeres do not. Mol. Ecol. 28, 3669–3680. (doi:10.1111/mec.15181)

26. Cawthon RM. 2002 Telomere measurement by quantitative PCR. Nucleic Acids Res. 30, e47. (doi:10.1093/nar/30.10.e47)

27. Bates D, Maechler M, Bolker BM, Walker SC. 2015 Fitting Linear Mixed-Effects Models Using lme4. J. Stat. Softw. 67, 1–48. (doi:10.18637/jss.v067.i01)

28. R Core Team. 2019 R: A Language and Environment for Statistical Computing. R Foundation for Statistical Computing. Vienna, Austria. https://www.R-project.org

29. van de Pol M, Wright J. 2009 A simple method for distinguishing within-versus between-subject effects using mixed models. Anim. Behav. 77, 753–758. (doi:10.1016/j.anbehav.2008.11.006)

30. Richards SA, Whittingham MJ, Stephens PA. 2011 Model selection and model averaging in behavioural ecology: the utility of the IT-AIC framework. Behav. Ecol. Sociobiol. 65, 77–89. (doi:10.1007/s00265-010-1035-8)

31. Urvik J, Rattiste K, Giraudeau M, Okuliarova M, Horak P, Sepp T. 2018 Age-specific patterns of maternal investment in common gull egg yolk. Biol. Lett. 14. 20180346. (doi:10.1098/rsbl.2018.0346)

32. Heidinger BJ, Herborn KA, Granroth-Wilding HMV, Boner W, Burthe S, Newell M, Wanless S, Daunt F, Monaghan P. 2016 Parental age influences offspring telomere loss. Funct. Ecol. 30, 1531–1538. (doi:10.1111/1365-2435.12630)

33. Lv L, Komdeur J, Li J, Scheiber IBR, Zhang Z. 2016 Breeding experience, but not mate retention, determines the breeding performance in a passerine bird. Behav. Ecol. 27, 1255–1262. (doi:10.1093/beheco/arw046)

34. Schroeder J, Nakagawa S, Rees M, Mannarelli M-E, Burke T. 2015 Reduced fitness in progeny from old parents in a natural population. Proc. Natl. Acad. Sci. U. S. A. 112, 4021–4025. (doi:10.1073/pnas.1422715112)

35. Noguera JC, Metcalfe NB, Monaghan P. 2018 Experimental demonstration that offspring fathered by old males have shorter telomeres and reduced lifespans. Proc. R. Soc. B 285, 20180268. (doi:10.1098/rspb.2018.0268)

36. Kroeger SB, Blumstein DT, Armitage KB, Reid JM, Martin JGA. 2020 Older mothers produce more successful daughters. Proc. Natl. Acad. Sci. U. S. A. 117, 4809–4814. (doi:10.1073/pnas.1908551117)

37. Brown AM, Wood EM, Capilla-Lasheras P, Harrison XA, Young AJ. 2020 Data from: Older mothers produce offspring with longer telomeres: a longitudinal within-parent analysis. Dryad Digit. Repository. (doi:10.5061/dryad.2jm63xsng)

