## Supplementary material for "Older mothers produce offspring with longer telomeres: a longitudinal within-parent analysis": Electronic Supplementary Material

### 1. Material and methods

*Study species and site*

Our study population comprises approximately 40 cooperative groups, each defending year-round territories in an approximately 1.5km$^2$ area of Tswalu Kalahari Reserve, South Africa. Individuals are identifiable in the field by their uniquely numbered metal ring and combination of three colour rings (SAFRING licence 1444). Adult individuals of the subspecies found at Tswalu (*P. mahali mahali*) have a conspicuous sex difference in beak colour and thus can be readily sexed in the field [1], while those that do not survive to adulthood are sexed using molecular techniques [2].

Dominant birds were identified from weekly observations of social groups. Dominant birds of both sexes show aggression (e.g. chasing, displacing) towards other group members more frequently, the dominant pair regularly sing together in a synchronised duet, and dominant males produce a solo song at dawn [1,3,4]. In the analyses presented in the main paper, the group's dominant breeding pair at the time of offspring laying were considered to be the parents of the offspring (for the purposes of calculating parental age), as previous genetic analyses have revealed that all eggs are laid by the dominant female and ~85% of offspring are sired by the resident dominant male (the remaining being sired by dominant males in other groups) [1]. Thus, the paternal age results in the main paper reflect the effects of the age of the offspring's social father (who contributes to their rearing), but this male is not always their genetic father. To establish whether these results hold (in particular paternal Δ age being a weaker predictor of offspring telomere length [TL] than maternal Δ age) when analysing the effects of paternal age for the *genetic* father (in case a paternal age effect is driven by gamete-mediated mechanisms for example) we then repeated the analysis using a smaller sample size of offspring for which microsatellite data were available and confirmed that the resident dominant male was also the genetic father of the offspring (see below for methods and outcome).

*Offspring and parental age calculations*

We included only known-age offspring in our analyses (i.e. birds that were first encountered as nestlings) to allow us to control for offspring age at sampling on RTL. Of the 356 offspring represented in our dataset, we knew the hatch date (and therefore age at sampling) of 316 offspring (89%) accurately to within one day, 345 accurately to within 3 days (98%) and all 356 (100%) accurately to within 12 days. We estimated the offspring's 'parental age' on the lay date of their clutch (modal clutch size = 2) by averaging the hatch date of all nestlings within the clutch and subtracting 16 days (the mean incubation period length). Parental ages were calculated as either the parent's known age (i.e. time since hatching, if the parent was first observed as a nestling) or their 'minimum age' (if the parent was first observed as a fledgling or an adult). If a parent was first seen as a fledgling, they were assumed to be a minimum of 30 days old when first seen. If a parent was first seen as an adult, they were assumed to be a minimum of 1 year old when first seen. Out of 60 fathers, 13 were first observed as nestlings, 8 as fledglings and 39 as adults. Out of 61 mothers, 22 were first observed as nestlings, 5 as fledglings, and 34 as adults. While estimating parental age in this way leads to some uncertainty in absolute parental age, once variation in parental age is partitioned into its within- and among-parent components (by within-individual mean centring [5]) this uncertainty passes entirely to the parental mean age component (which captures among-parent differences in mean age at conception), leaving accurate the parental Δ age component (the principal predictor of interest, as it captures within-parent differences in age at conception relative to that parent's mean parental age). Thus our findings regarding the effects of maternal Δ age and paternal Δ age can be interpreted with confidence, while some caution is needed with the interpretation of the effects of mean maternal and paternal age (from which one might otherwise draw inference about the extent of selective disappearance, for example [6]). Therefore, while we fit the mean age components in our full models alongside the Δ age components when partitioning the parental age effects, we do not draw inferences from our models about the strength of evidence for mean age effects.

*Blood sampling*

A small blood sample was collected from individuals of ≥3 days old and up to twice annually throughout their lives. The typical sampling regime involved taking small blood samples from each nestling in a clutch at 5, 13, and 30 days since the first nestling hatched, and most adult birds in the study population were also sampled at the start and end of each breeding season, yielding repeated samples throughout the lives of large numbers of individuals; both parents and their offspring. To collect blood from fledgling and adult birds, individuals were flushed from their roost chambers at night using custom-made capture bags. Blood was collected from the wing vein using a 26-guage needle and a non-heparinized microhaematocrit capillary tube, and stored at ambient temperature in ~500 µl of >99% ethanol until DNA extraction. DNA was extracted between May and August 2015 and stored in elution buffer at -20˚C until qPCR analysis (conducted between August 2015 and September 2016). To control for potential variation in telomere length estimates attributable to variation in the sampling date, we used a "sampling period" random effect, which identified the breeding season in which the sample was taken and whether it was taken early (September to December) or late (January to May) in that breeding season.

*Telomere length measurements*

The following methods have been described previously by Wood and Young 2019 [7]. DNA was extracted from blood samples using Gentra PureGene Genomic Purification Kits (Qiagen). Samples with poor DNA integrity (assessed using gel electrophoresis) were not included in analyses. DNA concentration and purity were assessed using a NanoVue 4282 Spectrophotometer (v1.7.3) with accepted ratios of between 1.7 and 2.0 for 260/280 and between 1.9 and 2.2 for 260/230. Whole blood telomere lengths were determined using quantitative PCR (qPCR) [8] with primers specific to *Plocepasser mahali* (GAPDH-F 5′-AAACCAGCCAAGTATGATGACAT–3′; GAPDH-R 5′-CCATCAGCAGCAGCCTTCA-3′; Tel1b 5′-CGGTTTGTTTGGGTTTGGGTTTGGGTTTGGGTTTGGGTT-3′; Tel2b 5′-GGCTTGCCTTACCCTTACCCTTACCCTTACCCTTACCCT-3′). GAPDH and telomere reactions were carried out on separate 96-well plates on a Stratagene Mx3000 instrument and each 20µl reaction contained 5ng DNA and 10µl SybrGreen fluorescent dye with low ROX (Agilent Technologies), with primers at a concentration of 200nM. The thermal cycling schedule for telomere reactions was 15 minutes at 95˚C, followed by 40 cycles of 95˚C for 15 seconds, 57˚C for 30 seconds, and 73˚C for 30 seconds. GAPDH reaction thermal cycles were the same, except the annealing temperature was 60˚C.

Each sample was run in triplicate for both GAPDH and telomere reactions. A calibration sample (containing DNA pooled from 3 individual birds) and a no-template control were also run in triplicate on every plate. A standard curve was created from the 5ng dilution of the calibration sample for each plate and efficiencies of GAPDH and telomere reactions were assessed using a 2x serial dilution (from 10ng to 0.625ng) of the calibration sample on every plate. For GAPDH plates, the mean standard curve efficiency was 97.91 (sd = 4.05, range = 90.7-109.4) with a mean $r^2$ of 1.00 (sd =

0.003, range = 0.988-1.000). For telomere plates, the mean standard curve efficiency was 98.65 (sd = 6.11, range = 85.80-112.80), with a mean $r^2$ of 1.00 (sd = 0.003, range = 0.990-1.000). Mean individual reaction efficiencies were 1.915 for GAPDH reactions (sd = 0.082; range = 1.654 – 2.258) and 1.80 for telomere reactions (sd = 0.060; range = 1.560–2.031). Mean reaction $r^2$ was 0.999 for both GAPDH and telomere reactions (GAPDH and telomere sd = 0.001; GAPDH range = 0.999-1.001, telomere range = 0.991-1.000).

In order to increase our ability to distinguish variation among plates from variation among birds, samples from birds from which more than one blood sample was taken were often split across plates to avoid having plates with a low number of birds (258 birds on 1 plate, 73 birds on 2 plates, 19 birds on 3 plates, 5 birds on 4 plates, and 1 bird on 5 plates). Although we attempted to run all samples in triplicate, any replicates with poor amplification (determined by its Cq value) were removed and mean RTL for each sample was calculated from the remaining replicates with good amplification. We calculated intra-class correlation coefficients (ICC) to estimate the inter- and intra- plate repeatability of technical replicates using the ‘consistency’ *icc* function in R (using the *irr* package [9]). The inter-plate repeatability for samples run on three different plates was 0.567 (two-way model with ‘single units’; 95% confidence interval [CI] = 0.339 - 0.772, n = 23 samples). We estimated the intra-plate repeatability of samples used in this study using a one-way model with average units. The ICC for GAPDH reaction triplicates was 0.986 (n = 729, 95% CI = 0.984 - 0.988). For GAPDH reaction duplicates (where a replicate was discarded due to poor amplification) the ICC was 0.988 (n = 36, 95% CI = 0.977 - 0.994). The ICC of telomere triplicates was 0.987 (n = 738, 95% CI = 0.986 - 0.989) and 0.973 for duplicates (n = 27, 95% CI = 0.941 - 0.988).

Relative telomere length (RTL) was estimated as the amount of telomere sequence per sample relative to a non-variable copy-number control gene, glyceraldehyde-3-phosphate-dehydrogenase (GAPDH) (following [10]). The following equation was used to calculate RTL, where $E_{TEL}$ and $E_{GAPDH}$ are the mean plate well efficiencies for telomeres and GAPDH respectively.

$$RTL = (E_{TEL} \wedge (Cq_{TEL[Calibrator]} - Cq_{TEL[Sample]})) / (E_{GAPDH} \wedge (Cq_{GAPDH[Calibrator]} - Cq_{GAPDH\ [Sample]}))$$

*Testing for the effects of paternal age for the genetic father*

The paternal age results in main paper reflect those for the offspring’s social father, who is typically also their genetic father. However, as ~15% of offspring are sired by males in other groups [1], and paternal age effects have the potential to be driven by the age of the *genetic* father too (e.g. via gamete-mediated mechanisms) we then established whether our findings hold when repeating the analysis using a smaller sample size of offspring for whom microsatellite data confirmed the resident dominant male was also their *genetic* father (see below for outcome). Microsatellite genotypes for 482 individuals at 13 markers were available from a previous study of our population (see [1] for genotyping protocols and multiplex design). We updated this data set by genotyping 928 additional individuals at these same 13 markers as well as an additional marker GCSW35 ([11]; which was included in multiplex 2 – see Table 1 in [1]). DNA for the latter 928 individuals was extracted from blood samples (preserved in >99% ethanol) using Qiagen DNeasy Blood & Tissue Kit (Qiagen), following manufacturer's instructions. PCRs were carried out in 2-µl reactions using Qiagen PCR Mastermix (Qiagen) at the following temperature profiles: 95°C for 15 min, followed by 35 cycles of 94°C for 30s, 56°C for 90s, and 72°C for 90s, with a final step of 60°C for 30 min. Samples were then genotyped on an ABI 3130xl Capillary Sequencer (Applied Biosystems, USA) and allele sizes scored and individually checked using Genemapper v4.0. To confirm comparable results between the first run of genotyping data (presented in [1]) and the newly genotyped individuals for this study, 125 samples were genotyped as part of both genotyping exercises. Comparison of genotypes for these 125 samples confirmed matching genotypes and allowed us to align and combine both data sets. The resident dominant male was deemed to be the genetic father whenever 1 or more alleles at every locus in the offspring matched one or more alleles at those same loci in the resident dominant male. Although not all 14 available loci were successfully genotyped for all individual birds, the minimum number of matching loci between an offspring-sire pair for which we inferred paternity was 7. This resulted in a data set of 533 telomere length measures from 258 offspring (with confirmed paternity) from 194 clutches born to 61 mothers and 57 fathers for use in the analysis (see 'Supporting analyses' below for the outcome).

*Statistical analysis*

Global models were fitted using the *lmer* function in the *lme4* package in R v.3.6.1 [12], and were fit using maximum likelihood. Models were fitted using the *bobyqa* optimizer to improve model convergence. Collinearity of variables was determined by assessing their variance inflation factors (VIFs) with the *car* package [13] as well as Pearson's correlation tests. Continuous fixed effect predictors were centred and scaled. Models with different fixed effect structures were ranked according to their AICc values using the *dredge* function in the *MuMIn* package [14]. The 'model nesting rule' was applied, which removes models from the top model set where a simpler nested version of that model was present with a lower AICc value (i.e. attracting stronger statistical support) [15,16].

### 2. Results

**Table S1.** Random effect estimates from top-ranked models presented in the main paper for (a) the population-level analysis and (b) the within-parent analysis. All random effects were fitted as random intercepts.

| Random effect | Variance |
|---|---|
| *(a) population-level analysis* | |
| Bird ID | 0.006119 |
| Mother ID/clutch ID | 0.002461 |
| Mother ID | 0.010233 |
| qPCR plate ID | 0.008166 |
| Sampling period | 0.003308 |
| *Residual* | 0.017960 |
| *(b) within-parent analysis* | |
| Bird ID | 0.006240 |
| Mother ID/clutch ID | 0.002347 |
| Mother ID | 0.009990 |
| qPCR plate ID | 0.008126 |
| Sampling period | 0.003358 |
| *Residual* | 0.017944 |

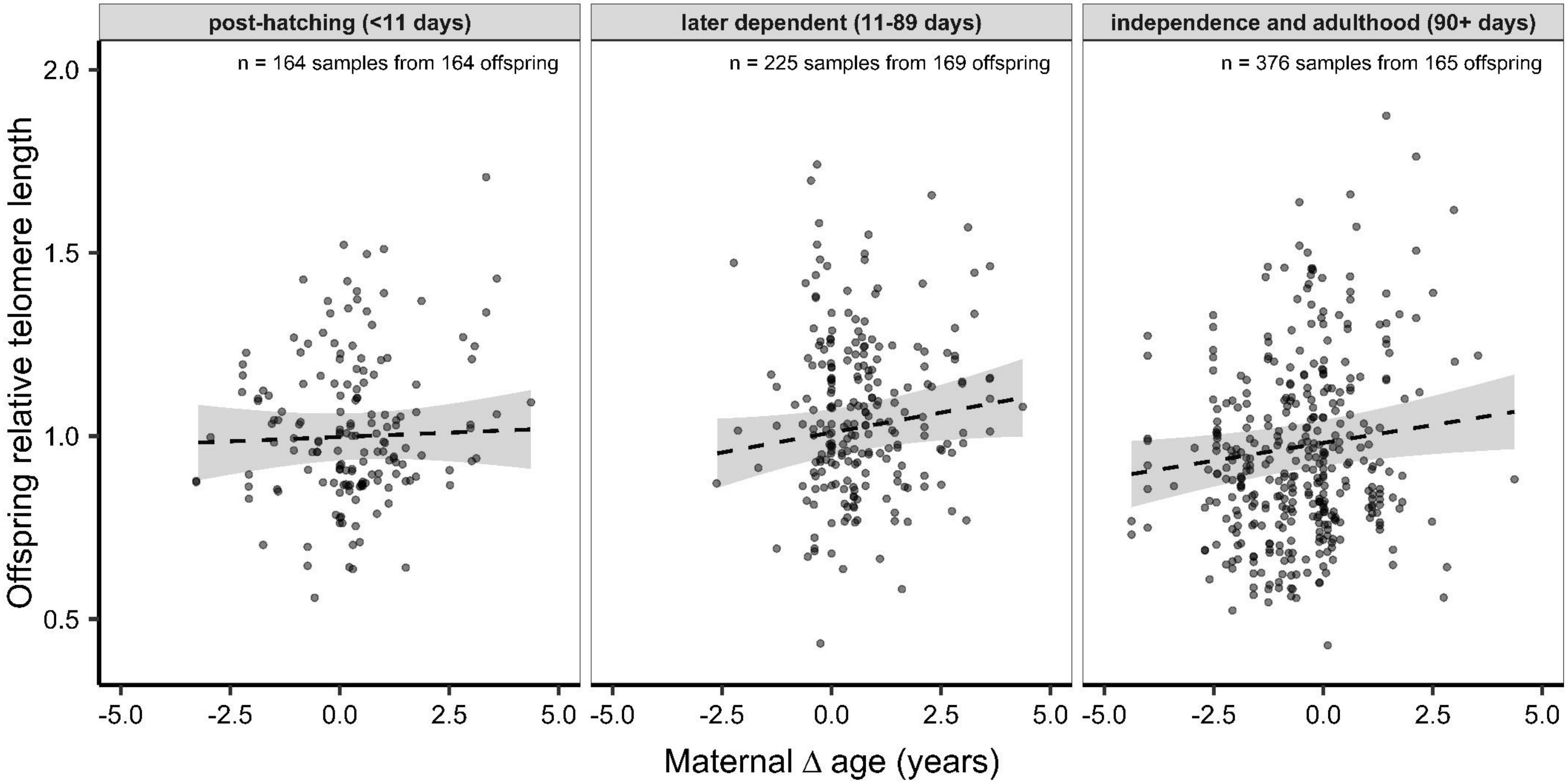


**Figure S1.** Relationship between maternal Δ age and offspring RTL for telomere length measurements taken at different stages through offspring development. Lines represent predictions from top-ranked model that included an interaction between maternal Δ age and offspring age class, which ranked at ΔAICc = 4.36 but did not appear in the top model set (Table 2 main paper) following removal of nested models. Effect sizes ± SE: *post-hatching period* = 0.007 ± 0.016, *later dependent period* = 0.031 ± 0.016, *independence and adulthood* = 0.028 ± 0.012.

### 3. Supporting analyses

*Our findings remain unchanged when focussing on the paternal age of the genetic father*

We repeated the key analysis presented in the main paper, using a smaller sample size of offspring for which microsatellite data were available and confirmed that the resident dominant male was their genetic as well as social father (see above). Genetic data have previously confirmed that the dominant female is always the mother [1]. Using this restricted data set of known maternity *and* paternity, the top model set was very similar to that presented in main paper: the top-ranked model contained only a positive effect of maternal Δ age (effect size ± S.E. = 0.031 ± 0.011), while a model containing only the effect of paternal Δ age was ranked second (ΔAICc = +1.52 relative to the top model; effect size ± S.E. = 0.030 ± 0.012). The null model, containing no parental Δ age effects (maternal or paternal), was ranked third (ΔAICc = +5.42 relative to the top model). The random effects structure in the global model used here differed slightly from that presented in the main paper: while we retained the mother ID random effect, we removed clutch ID as its inclusion (when nested within mother ID as per the analysis in the main paper) was resulting in the over-fitting of this model (the nested random effect estimated almost zero variance). Unlike the analysis on the full dataset presented in the main paper, we were able to fit both mother and father ID without producing singular fits (after removing clutch ID), and therefore included both variables as random intercepts.

*Our findings cannot be attributed to variation in social group size or helper number*

Sparrow-weaver groups comprise a dominant breeding pair and zero to 12 non-breeding subordinate helpers of both sexes that assist with feeding young [1]. The detected positive effect of maternal age on offspring telomere length cannot be attributed instead to correlated variation in social group size or helper number (group size minus 2) as (i) maternal age and social group size at offspring rearing (mean number of adults >6 months old in the offspring's group, when the offspring was 1-30 days of age) are not significantly correlated (Pearson's: $r = -0.06$, $t = -1.54$, $df = 746$, $p = 0.12$; $n = 349$ offspring from 241 breeding attempts by 61 mothers), and (ii) including the offspring's social group size at rearing as an additional predictor in the global model presented in the main paper leaves our findings qualitatively unchanged: maternal Δ age (and hence not paternal Δ age) remains in the top model and third-ranked models down to ΔAICc = +2.57 (maternal Δ age effect size ± SE in the best-supported model = 0.030 ± 0.009, n = 748 telomere length measures from 349 offspring for which rearing group size information was available). The second-ranked model (which contained paternal but not maternal Δ age) had a ΔAICc = +2.10.

*The maternal age effect on offspring TL appears to persist into adulthood*

To investigate whether the positive maternal age effect on offspring TL detected in the main paper persists into the offspring's adulthood, we repeated the key analysis presented in the main paper, but this time factorising offspring age into two classes: <1 year old and >1 year old. The model containing maternal Δ age, offspring age class, and the interaction between them (which did not appear in the top model set after removal of nested models) revealed that the maternal Δ age effect on offspring TL is present in adult offspring (maternal Δ age effect size in >1 year old offspring ± SE = 0.035 ± 0.013; maternal Δ age effect size in <1 year old offspring ± SE = 0.013 ± 0.013).

### References


1. Harrison XA, York JE, Cram DL, Hares MC, Young AJ. 2013 Complete reproductive skew within white-browed sparrow weaver groups despite outbreeding opportunities for subordinates of both sexes. *Behav. Ecol. Sociobiol.* **67**, 1915–1929. (doi:10.1007/s00265-013-1599-1)

2. Wood EM. 2017 Causes and fitness consequences of telomere dynamics in a wild social bird. PhD thesis, University of Exeter, Penryn, Cornwall.

3. Harrison XA, York JE, Cram DL, Young AJ. 2013 Extra-group mating increases inbreeding risk in a cooperatively breeding bird. *Mol. Ecol.* **22**, 5700–5715. (doi:10.1111/mec.12505)

4. York JE, Radford AN, de Vries B, Groothuis TG, Young AJ. 2016 Dominance-related seasonal song production is unrelated to circulating testosterone in a subtropical songbird. *Gen. Comp. Endocrinol.* **233**, 43–52. (doi:10.1016/j.ygcen.2016.05.011)

5. van de Pol M, Wright J. 2009 A simple method for distinguishing within- versus between-subject effects using mixed models. *Anim. Behav.* **77**, 753–758. (doi:10.1016/j.anbehav.2008.11.006)

6. Beirne C, Delahay R, Hares M, Young A. 2014 Age-related declines and disease-associated variation in immune cell telomere length in a wild mammal. *PLoS One* **9**, 1–6. (doi:10.1371/journal.pone.0108964)

7. Wood EM, Young AJ. 2019 Telomere attrition predicts reduced survival in a wild social bird, but short telomeres do not. *Mol. Ecol.* **28**, 3669–3680. (doi:10.1111/mec.15181)

8. Cawthon RM. 2002 Telomere measurement by quantitative PCR. *Nucleic Acids Res.* **30**, e47. (doi:10.1093/nar/30.10.e47)

9. Gamer M, Lemon J, Singh P. 2019 irr: Various coefficients of interrater reliability and agreement. R package version 0.84.1. hhttps://CRAN.R-project.org/package=irr.

10. Pfaffl MW. 2001 A new mathematical model for relative quantification in real-time RT–PCR. *Nucleic Acids Res.* **29**, e45. (doi:10.1111/j.1365-2966.2012.21196.x)

11. Mcrae SB, Emlen ST, Rubenstein DR, Bogdanowicz SM. 2005 Polymorphic microsatellite loci in a plural breeder, the grey-capped social weaver (*Pseudonigrita arnaudi*), isolated with an improved enrichment protocol using fragment size-selection. *Mol. Ecol. Notes* **5**, 16–20. (doi:10.1111/j.1471-8286.2004.00816.x)

12. R Core Team. 2019 R: A Language and Environment for Statistical Computing. R Foundation for Statistical Computing. Vienna, Austria. https://www.R-project.org

13. Fox J., Weisberg S. 2019 An R Companion to Applied Regression. Third edition. Sage, Thousand Oaks CA. https://socialsciences.mcmaster.ca/jfox/Books/Companion/.

14. Barton K. 2018 MuMIn: Multi-Model Inference. R Package version 1.43.6.

15. Richards SA, Whittingham MJ, Stephens PA. 2011 Model selection and model averaging in behavioural ecology: the utility of the IT-AIC framework. *Behav. Ecol. Sociobiol.* **65**, 77–89. (doi:10.1007/s00265-010-1035-8)

16. Harrison XA, Donaldson L, Correa-Cano ME, Evans J, Fisher DN, Goodwin CED, Robinson BS, Hodgson DJ, Inger R. 2018 A brief introduction to mixed effects modelling and multi-model inference in ecology. *PeerJ* **2018**, e4794. (doi:10.7717/peerj.4794)
